# Ketones facilitate transcriptional resolution of secondary DNA structures in premature aging

**DOI:** 10.1101/2022.03.08.483430

**Authors:** Michael Angelo Petr, Lina M Carmona-Marin, Tulika Tulika, Stella Kristensen, Simon Reves, Daniela Bakula, Guido Keijzers, Brenna Osborne, Sarah J Mitchell, Sam Hamilton, Jonathan Kato, Irene Alfaras, Amanuel A. Teklu, Indra Heckenbach, Jakob Madsen, Michael Ben Ezra, Garik Mkrtchyan, Erika Varner, Benjamin Fink, Eliana von Krusenstiern, Nathaniel W Snyder, Hector Herranz, Rafael de Cabo, Morten Scheibye-Knudsen

**Affiliations:** Center for Healthy Aging, Department of Cellular and Molecular Medicine, University of Copenhagen, Copenhagen, Denmark; Translational Gerontology Branch, National Institute on Aging, NIH, Baltimore, MD 21224, USA; Tracked.bio, Copenhagen, Denmark; Department of Biotechnology and Biomedicine, Technical University of Denmark, Lyngby, Denmark; Department of Health Sciences and Technology, ETH Zurich, Zurich, Switzerland; Aging Institute, University of Pittsburgh/UPMC, Pittsburgh, PA 15219 USA; Department of Biochemistry and Molecular Biology, College of Health Sciences, Mekelle University, Ethiopia; Buck Institute for Research on Aging, Novato, CA 94945, USA; Center for Metabolic Disease Research, Department of Cardiovascular Sciences, Lewis Katz School of Medicine, Temple University, Philadelphia, PA 19140; Department of Cellular and Molecular Medicine, University of Copenhagen, Copenhagen, Denmark

**Keywords:** Aging, Cockayne syndrome, Ketogenic diet, Secondary DNA structures, G-quadruplex

## Abstract

There is currently no established intervention for Cockayne syndrome, a disease characterized by progressive early onset neurodegeneration with features of premature aging. Here, we tested if acetyl-CoA precursors, citrate and beta-hydroxybutyrate, could reduce features of Cockayne syndrome in three model systems. We identified the gene Helicase 89B as a homologue of CSB in drosophila and found that the ketone beta-hydroxybutyrate rescued features of premature aging in Hel89B deficient flies. In mammals, loss of the citrate carrier Indy exacerbated the phenotype of Csb^m/m^ mice which was rescued by a ketogenic diet. The rescue effect appeared to be mediated through ketone stimulated histone acetylation and facilitation of transcriptional readthrough of secondary DNA structures. These findings link a ketogenic diet with transcriptional resolution of secondary structures and DNA repair.

## Introduction

Cockayne syndrome (CS) is a rare genetic disease characterized by neurological dysfunction, photosensitivity and premature aging^1,2,3^. The disease is most commonly caused by mutations in the CSB (ERCC6) or CSA (ERCC8) genes, two genes involved in transcription and transcription coupled nucleotide excision DNA repair. Previously, it has been shown that acetyl-CoA levels are decreased in models of Cockayne syndrome likely through persistent activation of a poly-ADP-ribose polymerase 1 (PARP1) mediated DNA damage response, loss of Nicotinamide adenine dinucleotide (NAD) and shunting of pyruvate to lactate^4^. This could likely contribute to a decrease in acetyl-CoA dependent phenotypes such as demyelination and neurological dysfunction. Citrate and ketones intimately regulate the amounts of acetyl-CoA. Notably, a ketogenic diet (KD) is neuroprotective in mice and increases the lifespan in nematodes^4^ and mice^5,6^. Interestingly, acetyl-CoA regulates the amount of histone acetylation and histone deacetylation^7,8,9,10^ which are mediated via histone acetyltransferases (HATs) and histone deacetylases (HDACs), respectively. Acetyl-CoA acts as the acyl-donor molecule for these acetylation reactions and metabolism could thereby directly impacts the chromatin landscape ^11^. To that end, there is an overall depletion of the necessary metabolites and CoA pools with age, affecting histones and transcription^12^. Without proper histone acetylation, especially in the context of premature aging this may lead to detrimental chromatin remodeling^13^.

Indy, a citrate transporter, was originally discovered in drosophila and heterozygote mutations in Indy significantly extended lifespan of *D. melanogaster* and *C. elegans*^14,15^. In mice, the mIndy^-/-^ (Slc13a5) mouse has decreased adiposity and increased insulin resistance, features also reported in Csb^m/m^ mice^16^. Interestingly, patients with SLC13A5 mutations suffer from early onset epilepsy, neurodegeneration and developmental delay ^17,18^ overlapping with features seen in Cockayne syndrome ^1,19^. We hypothesized that loss of Indy would exacerbate Cockayne syndrome phenotypes and that we could rescue this exacerbation by adding ketones to titrate acetyl-CoA back. To test this, we investigate how citrate and ketone metabolism might impact Cockayne syndrome and aging phenotypes in drosophila, mice and human cell lines. Indeed, we discovered the existence of transcription coupled repair of DNA in drosophila via the CSB homologue Hel89B and found that ketones can modulate DNA repair and extend lifespan in drosophila. To test this in mammals, we generated a novel Csb^m/m^/mIndy^-/-^ double knockout (DKO) mouse and observed a similar outcome. Mechanistically, ketones appear to facilitate the resolution of transcription through secondary structures by increasing H3K27 acetylation. In short, we have discovered a possible mechanism for the neuroprotective effect of ketones.

## Results

### Hel89B is a drosophila CSB homologue

To understand if a ketogenic diet could impact the lifespan of CSB deficient animals we turned to *Drosophila melanogaster*, a model that has been used extensively for aging studies and where INDY is known to influence lifespan^14^. We performed hierarchical clustering of SWI/SNF like proteins in drosophila with human CSB and found the closest homologue of CSB in fruit flies appears to be Helicase 89, isoform B (Hel89B) (Figure 1A). We next generated Hel89B knockdown flies using the Gal4 system and observed that Hel89B-Gal4 flies exhibit a significantly reduced hatch rate compared to their controls (Figure 1B, Supplementary Figure 1A). A key phenotype of nucleotide excision repair defects in patients is sensitivity to UV-C irradiation. Indeed, UAS-Hel89B (Hel89B) larvae (Supplementary Figure 1B) exhibits reduced survival with increasing UV-C doses, compared to their wild-type counterparts UAS-mGFP (mGFP) (Figure 1C, D, Supplementary Figure 1C). Strikingly, supplementing larvae with 10mM ketones significantly improved the survival of both Hel89B and mGFP larvae after UV damage compared with untreated controls.

**Figure 1.**
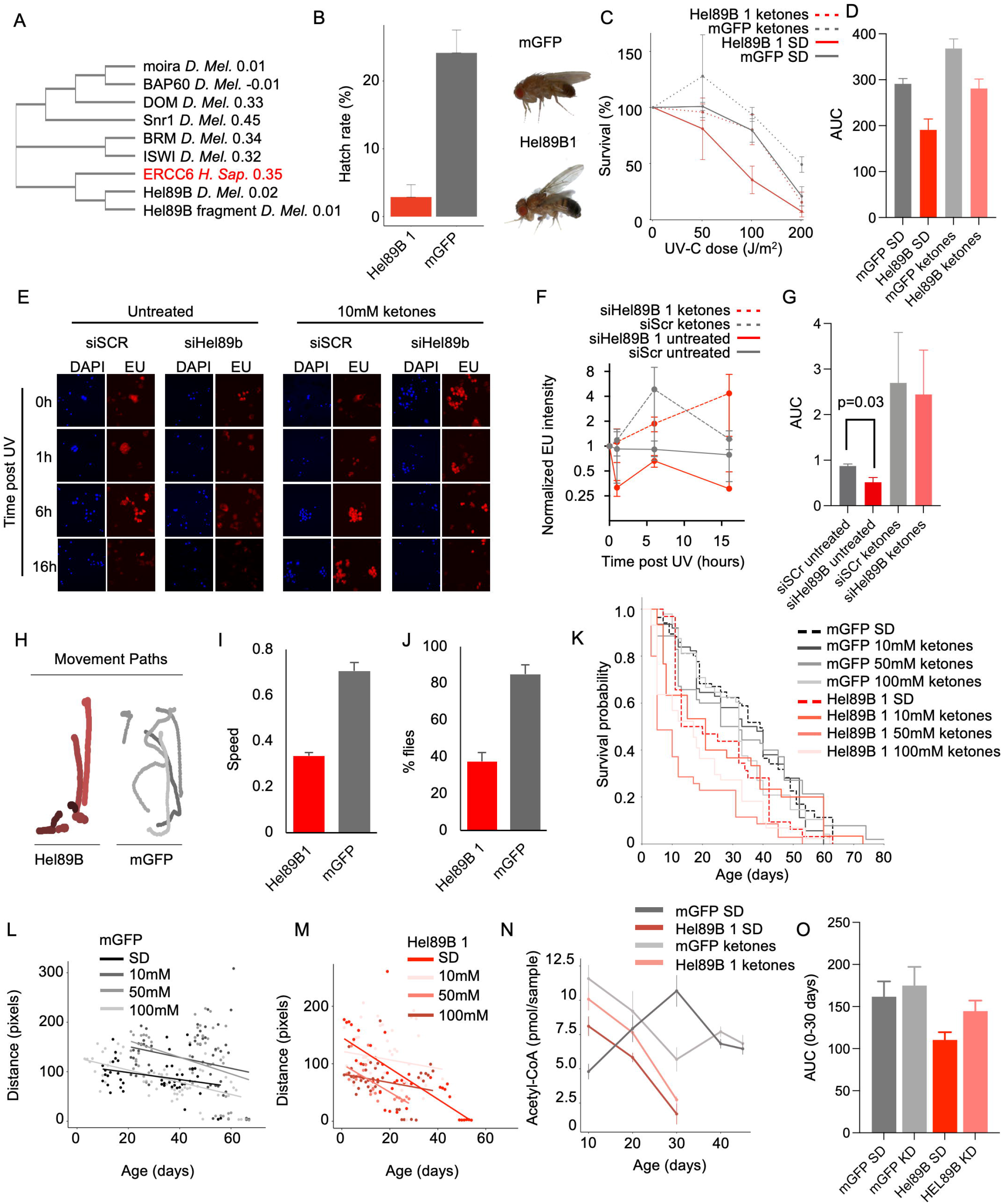
Hel89B is a Drosophila melanogaster homologue of CSB. **A** Hierarchical clustering of Drosophila SWI/SNF genes with human ERCC6 based on the sequence. **B** Hatch rate of Hel89B1 and mGFP flies (n = 6, 25-30 flies per replica). **C** Survival of standard or 10mM ketone fed larvae after exposure to different doses of UV-C light. **D** AUC of UV survival graphs in C. **E** Representative images of resumption of RNA synthesis after UV induced damage. **F** Quantification of EU staining from E (300-900 cells per replicate. N = 3). **G** AUC of F. **H** Negative geotaxis flowerplot of mGFP and Hel89B1 flies. **I** Speed from the negative geotaxis assay. **J** % of flies that cross 5cm line after 5 seconds from negative geotaxis assay. **K** Lifespan of flies treated with various doses of the ketone betahydroxybutyrate. **L** Max distance traveled per day of mGFP flies in K. **M** Max distance traveled per day of Hel89B flies in K. **N** Acetyl-CoA measurements of flies. **O** AUC of acetyl-CoA measurements from N.

A major hallmark of transcription coupled repair deficiency is a delayed resumption of RNA synthesis after UV-C irradiation. We therefore tested whether resumption of RNA synthesis was delayed upon knocking down Hel89B in Drosophila S2 cells (Supplementary Figure 1D). Indeed, these cells displayed decreased EU fluorescence within the cell nuclei, indicating reduced RNA synthesis with increasing time after exposure to UV light (Figure 1E). Quite strikingly, ketones facilitated faster resumption of RNA synthesis post UV exposure (Figure 1E,F,G). In sum, these results indicate that nucleotide excision repair exists in flies, that Hel89B may be a functional CSB homologue and that ketones somehow attenuate transcription coupled repair defects.

### Hel89B knockdown leads to impaired motor function in drosophila

A known phenotype in Cockayne syndrome is worsening of gait and motor function. We therefore wanted to assess this phenomenon in our Hel89B flies. We first turned to the negative geotaxis assay, with inspiration from the RING assay in flies ^20^, utilizing custom build hardware and software to track and quantify the behavior of the flies. The crawling profile (coined, flower plot) of the flies to reach the top after being displaced to the bottom of the vial was limited in complexity in the Hel89B flies compared to their mGFP controls (Figure 1H). The speed of the Hel89B flies was also significantly lower (Figure 1I), and the time it took for the Hel89B flies to vertically crawl 5cm in the vial was longer (Figure 1J).

To determine our treatment regimen for flies, we wanted to assess which dosage of ketones was optimal. We tested three doses of the ketone beta-hydroxy butyrate (BHB): 10mM, 50mM, and 100mM throughout the lifespan of Hel89B and control flies (Figure 1K). 10mM ketones appear to have a mild lifespan increase towards the end of life in both genotypes, while 50mM and 100mM BHB appear to be detrimental. Furthermore, by tracking the motor function of flies over time, we found that fly movement is reduced with age and that 10mM BHB treatment conserves motor function in both mGFP (Figure 1L) Hel89B flies (Figure 1M).

We next wanted to investigate whether ketones could increase the levels of acetyl-CoA in our Hel89B flies. We therefore measured acyl-CoAs using mass spectrometry across the lifespan in the flies. In the Hel89B flies, we observed an overall decrease in acetyl-CoA throughout their life and this was ameliorated by ketones (Figure 1N,O and Supplementary Figure 1E). In sum, these results indicate that Hel89B flies indeed have impaired motor function, that 10mM ketones is a potentially beneficial dose and that this dose increases acetyl-CoA levels in Hel89B flies.

### Ketones improves the motor function of flies

To understand the interaction between the PARP1-acetyl-CoA axis, INDY, and Hel89B a large number of experiments were needed. We therefore developed a high throughput life- and health-span platform using 3D printed parts, computer vision, and deep neural networks (Figure 2A). Specifically, we trained convolutional neural networks to identify flies in video feeds of their lab habitat in real time allowing us precise and high-throughput tracking of flies longitudinally. This allowed us to monitor the fruit flies continuously for the duration of each individual fly’s life and predict the age and health of the fly at any point. With this novel methodology, we characterized the lifespan and age-associated behavioral changes of a battery of genotypes, ketones, citrate and Olaparib (20 genotypes and 4 treatment groups, 5644 flies) (Supplementary Figure 2). As confirmed before^21^, heterozygote INDY flies live longer than the homozygote counterparts and wild-type flies (Figure 2B, Figure 2C) and INDY206/Hel89B1 mutants, who are relatively short-lived, have increased max and mean lifespans (Figure 2B, C) compared to Hel89B mutants. We next quantified the motor function including distance traveled and walking speed for all of our flies using this system. We found that 10mM BHB improved the lifetime motor function (Figure 2D) and change in speed with age (Figure 2E) of most genotypes tested including Hel89B1 flies. Interestingly, this was also the case for citrate treatments that slightly increased mean lifespan in many genotypes. Interestingly, ketones overall increase the starting distance of flies (Figure 1F), however decrease the change of speed throughout their lifespan (Figure 1G). longitudinal results show that overall, ketones improve the lifespan of certain genotypes and appear to improve the lifetime motor function of flies.

**Figure 2.**
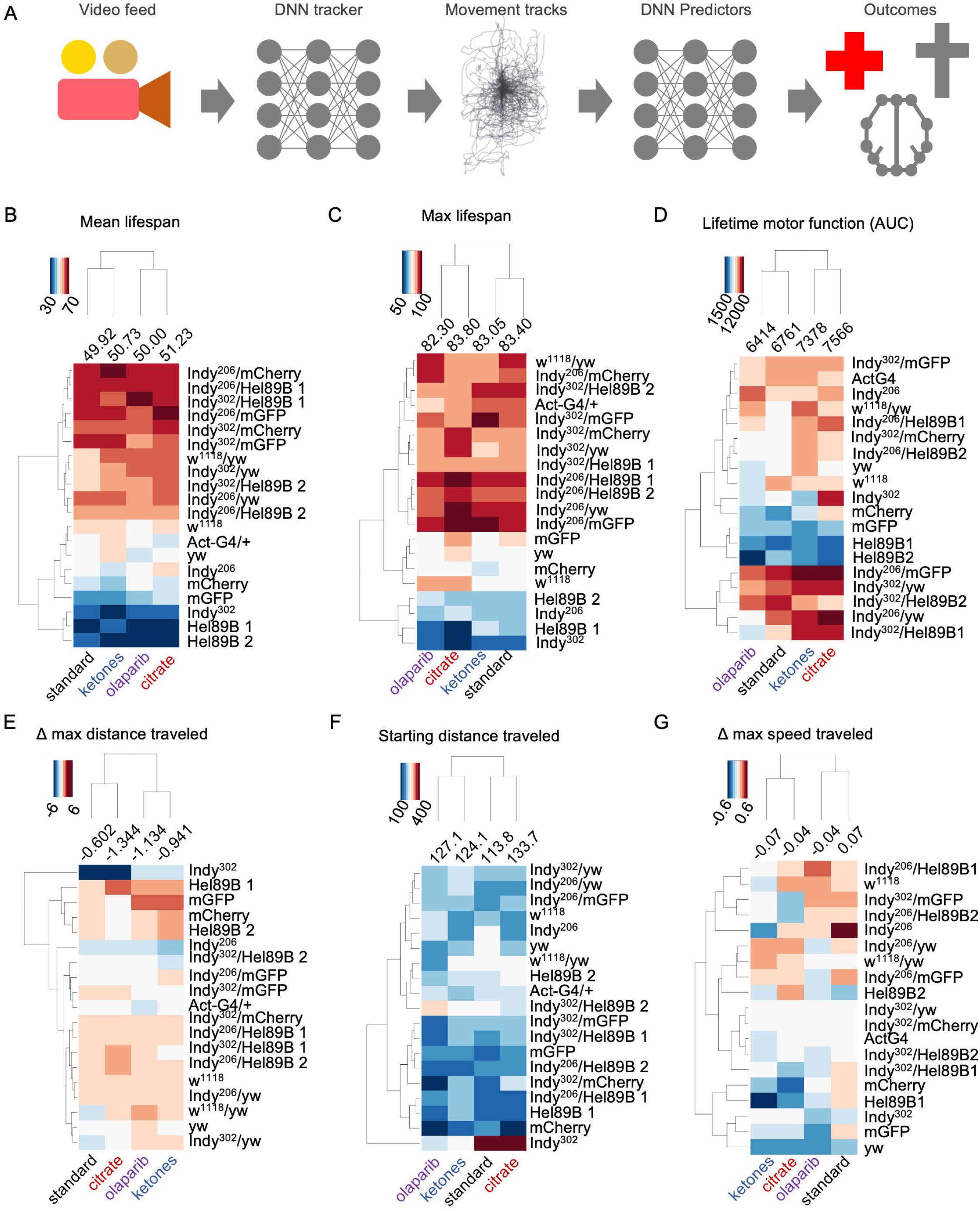
High-throughput phenotyping show ketones rescue motor function in normal and premature aging. **A** Analysis pipeline of experiments. **B** Heatmap of mean lifespan (n=3, 5644 flies). **C** Heatmap of max lifespan (n=3, 5644 flies). **D** Lifetime motor function: AUC of distance traveled until mean lifespan (n=3, 5644 flies). **E** Change in the max distance traveled as a function of age (n=3, 5644 flies). **F** Starting distance traveled (n=3, 5644 flies). **G** Change in the max speed traveled as a function of age (n=3, 5644 flies).

### A ketogenic diet rescues premature aging in Csb-mIndy double knockout mice

Since there are large variations in how different organisms handle acetyl-CoA, citrate and ketone metabolism, we next wanted to investigate mice. To understand if acetyl-CoA precursors could affect the phenotype of Csb^m/m^ we crossed the Csb^m/m^ mice with mIndy^-/-^ mice to generate Csb^m/m^ mIndy^-/-^ double knockout (DKO) mice. We then fed 6-month-old WT, Csb^m/m^, mIndy^-/-^ and DKO a standard (SD) or a ketogenic diet (KD) for 6 months (Supplementary Figure 3A). Throughout the study the mice were monitored with key metrics including body weight, food intake, body temperature, body composition and gait function (Figure 3A–C, Supplementary Figure 2E). As we hypothesized, the DKO mice exhibited exacerbated phenotypes, and benefited the most from the ketogenic diet. The DKO mice had lower body weight that was rescued by the ketogenic diet despite no difference in food intake or temperature (Figure 3A–D). The DKO mice also displayed an inability to accumulate fat, a characteristic of DNA repair deficiency, on a standard diet, however this was rescued when fed a ketogenic diet (Figure 3E–G). This was also confirmed when the mice were dissected and white adipose tissue and other organs were weighed (Supplementary Figure 3B). After observing the fat modulation in the DKO mice, we decided to analyze the *in vivo* metabolism of the mice with enclosed metabolic cages. Here, we observed that DKO mice had an overall higher oxygen consumption rate compared to wild type mice and this was greatly reduced by a ketogenic diet in the double knockout mice (Figure 3H). The respiratory exchange ratio (RER) revealed that Csb^m/m^ single knockouts switched to use of fats as an energy substrate (Figure 3I). We next investigated the metabolic output of the mice while they ran on a metabolic treadmill, to gauge energetic efficiency. Csb^m/m^ mice exhibited an overall lower VO2 (Figure 3J) and RER (Figure 3K), indicating utilization of fat as an energy source while active. When administered the ketogenic diet, the Csb^m/m^ mice have a comparable RER to the WT mice, however.

**Figure 3.**
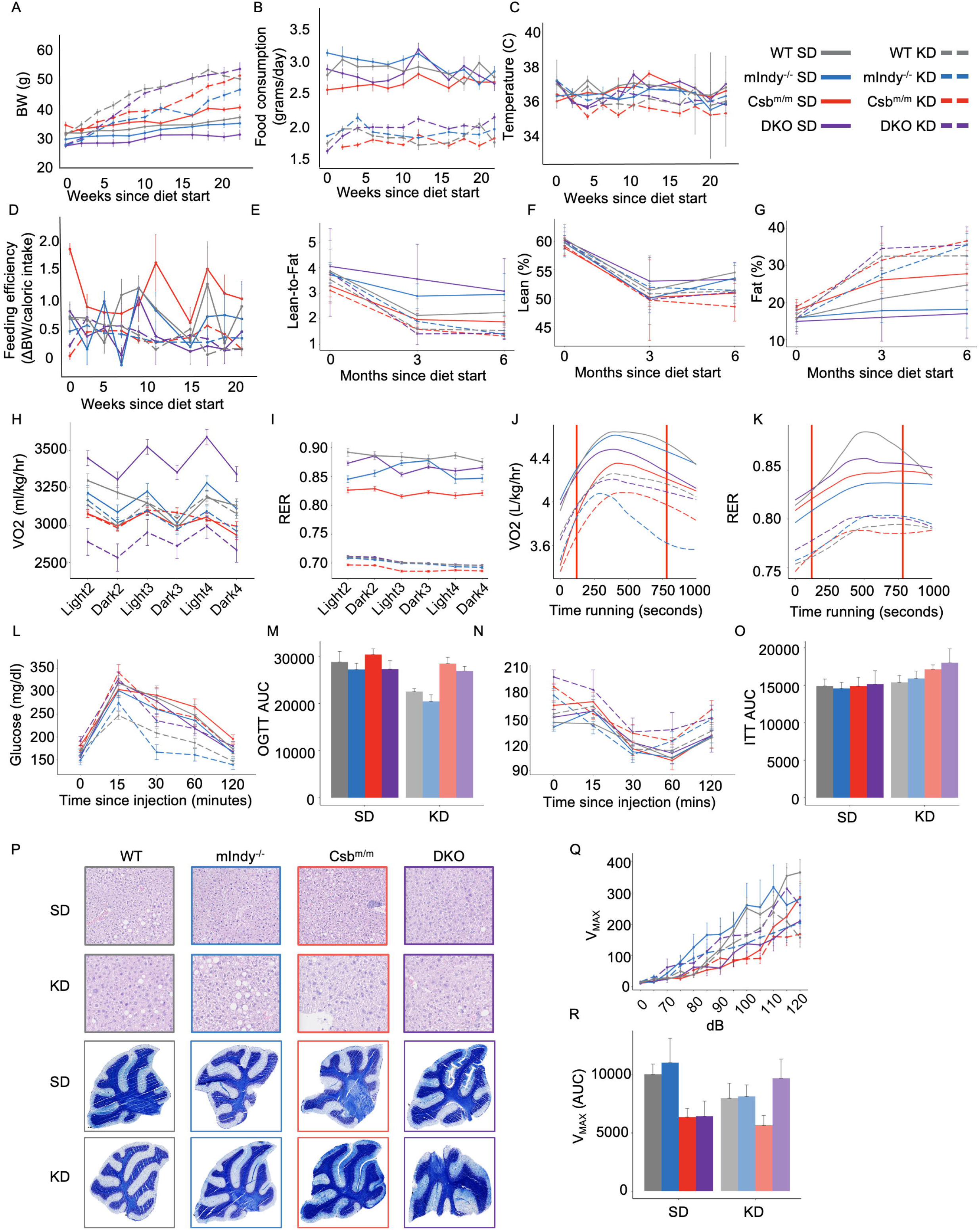
A ketogenic diet rescues features of exacerbated aging in Csb^m/m^ / mIndy^-/-^ mice. **A** Body weight of 6 month old (start) WT, mIndy^-/-^, Csb ^m/m^ and Csb^m/m^ / Indy^-/-^ double knockouts (DKO). **B** Food consumption. **C** Body temperature. **D** Feeding efficiency. **E** Lean-to-fat ratio. **F** Percentage lean mass. **G** Fat percentage. **H** VO_2_ of the mice in metabolic cages. **I** RER of the mice in metabolic cages. **J** VO_2_ of the mice in metabolic treadmills. **K** RER of the mice in metabolic treadmills. **L** Oral Glucose Tolerance Test (OGTT). **M** AUC of OGTT. **N** Insulin Tolerance Test (ITT). **O** AUC of ITT. **P** Representative micrographs of Haematoxylin and eosin (H&E) staining of liver tissues and luxol fast blue staining of cerebellum. **Q** Startle chamber analysis. Maximum voltage (Vmax) measured from the piezoelectric sensor at each decibel (dB) sound. **R** AUC of Q. n=12 per group, male C57BL/6J. Data presented with standard error of means (SEM).

In the past it has been reported that Csb^m/m^ mice are able to clear glucose at a higher rate than WT mice^3^. In our study, we found that the ketogenic diet improved glucose tolerance in WT and mIndy^-/-^ mice but not in Csb^m/m^ or DKO mice (Figure 3L,M). We observed little effect on insulin tolerance between the groups (Figure 3N,O). Further, hematoxylin and eosin (H&E) staining was performed on the postmortem liver, and decreased lipid droplet formation in the livers of the Csb^m/m^, mIndy^-/-^ and DKO mice was observed (Figure 3P). In all genotypes, when administered the ketogenic diet, these mice experienced an increase of fat droplets in the liver. Notably, luxol fast blue staining of the cerebellum revealed that Csb^m/m^ mice displayed a decreased amount of stained myelin, a feature previously seen in Cockayne syndrome patients ^23^, and this was reversed when given a KD increased (Figure 3P). We also investigated the hematology of the mice and found increased levels of large unstained cells and monocytes upon Csb deficiency perhaps suggesting increased inflammatory activation (Supplementary Figure 3C). In sum, loss of mIndy exacerbates the phenotype of Csb^m/m^ mice and a ketogenic diet attenuates these changes.

### A ketogenic diet attenuates neurological deficits in double knockout mice

We wanted to further assess the neurological function of the mice, given that CS patients experience neurological dysfunction, such as hearing loss^24,25,26,19^. Therefore, we utilized a piezoelectric-based startle chamber (SR Labs) for testing the hearing in the mice. Csb^m/m^ and DKO mice have hearing deficits, and furthermore the ketogenic diet improved the hearing in the DKO mice (Figure 3Q) by showing an increased sensitivity in the hearing test. Behavior attributes including open-field and elevated zero maze were used to assess changes in anxiety behavior and activity (Supplementary Figure 2D,E). Cockayne syndrome patients are described to be outgoing^25^, and Csb^m/m^ mice were previously observed as less anxious from open-field examination^4^. However, we did not observe the same in our mice in the open-field test. Y-maze spontaneous alteration test was also carried out to assess the overall cognitive function of the mice, to determine if working memory was improved by the ketogenic diet, however we saw no genotype or diet effects (Supplementary Figure 2F). Lastly, high resolution kinematic analysis was performed on the mice, specifically to measure gait speed which is a metric translatable to humans and declines with age^27^. Overall the knockout mice had an increased gait speed, and the ketogenic diet normalizes this (Supplementary Figure 2G).

### A ketogenic diet reduces transcriptomic changes in mIndy^-/-^, Csb^m/m^ and double knockout mice

To further evaluate global changes in the brain of the mouse models we performed RNA-seq analysis of the cerebellum, a target organ in Cockayne syndrome, in the four genotypes with and without a ketogenic diet (Supplementary Figure 4A). Notably, the KD appeared to reduce the number of significantly changed gene ontology pathways (Supplementary Figure 4B). Further, Csb^m/m^ and DKO mice have an upregulation in inflammatory pathways, consistent with previous findings ^24,23^, and the ketogenic diet suppresses these pathways (Figure 4A and Supplementary data file). In general, the ketogenic diet upregulated mitochondrial energy related pathways as well as pathways involved in ribosome function across genotypes. In both standard and ketogenic diets, the DKO mice have downregulated stimuli and smell related pathways. The olfactory down-regulation is observed in the Csb^m/m^ and mIndy^-/-^ mice as well (Figure 4A). Interestingly, mIndy^-/-^ mice on a ketogenic diet did not have a single significantly changed downregulated GO term and only a few upregulated suggesting normalization of the cerebellar transcriptome in this genotype.

**Figure 4.**
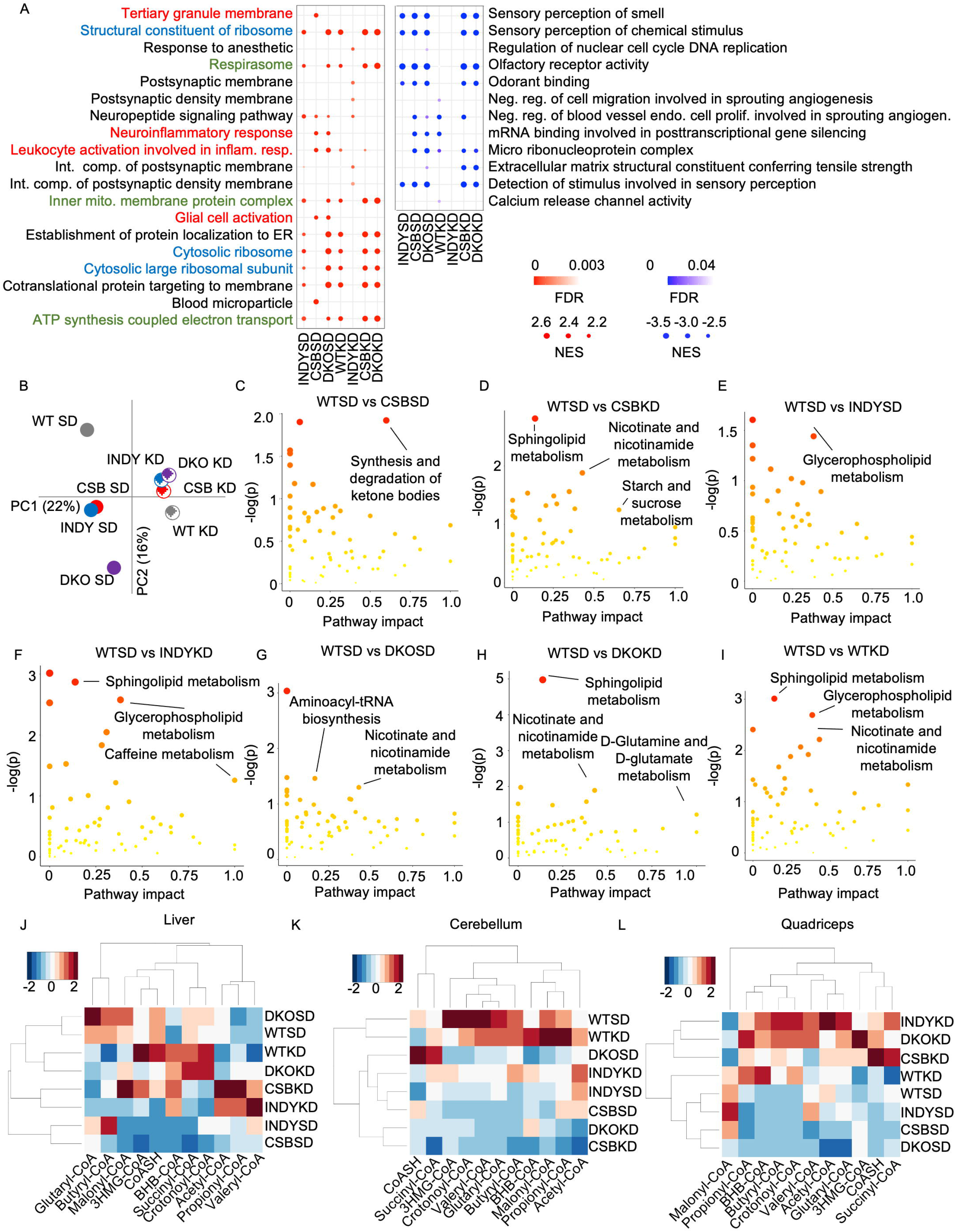
A ketogenic diet attenuates transcriptional and metabolic consequences of premature aging. **A** Top-5 altered Gene ontology (GO) terms from each genotype in RNA-sequencing of the cerebellum (n=6 per group). Red: up Blue: down (see supplementary data for the complete list of GO terms). **B** Principal component analysis of ~6000 metabolites in the cerebellum (n=6 per group). **C-I** Pathway topology analysis of the metabolomics experiments. All groups are compared to WTSD. **J** Acyl-CoA species heatmaps of liver, cerebellum (**K**) and quadriceps (**L**) (Z-score normalized within each metabolite, n=6).

### A ketogenic diet normalizes the metabolome of Csb^m/m^, mIndy^-/-^ and DKO mice

Since we observed global *in vivo* metabolic changes in the four genotypes, we decided to perform untargeted metabolomics on the cerebellum, a target organ in Cockayne syndrome. To get a broad overview of changes, we first performed hierarchical clustering and principal component analysis (Figure 4B and Supplementary Figure 4C). Strikingly, Csb^m/m^ and mIndy^-/-^ mice displayed almost identical metabolic profiles while the DKO further exacerbated the difference relative to WT. The ketogenic diet attenuated the genotype component (PC2) of the metabolic profile of mIndy^-/-^, Csb^m/m^ and DKO mice (Figure 4B). We further investigated specific pathways that were altered compared to WT mice fed standard food. In the enrichment analysis (Supplementary Figure 4), sphingolipid metabolism was greatly upregulated by the ketogenic diet, of note because sphingolipid metabolism is essential to myelin synthesis, which counteracts the demyelination phenotype in CS patients^28^. When considering the pathway topology impact compared to WT SD, one of the most significant pathways that appeared were “synthesis and degradation of ketone bodies” in Csb^m/m^ mice (Figure 4C), which could indicate that ketones metabolism is dysregulated in the Cockayne syndrome mice. Considering the pathway impact, nicotinate and nicotinamide are upregulated when the Csb^m/m^ mice are administered a KD (Figure 4D), driving the point that a KD may improve NAD^+^ metabolism. Of considerable interest as well, is the increased impact of aminoacyl-tRNA biosynthesis in the DKO SD group versus DKO KD (Figure 4G), perhaps caused by Csb’s role in transcription guided resolution of secondary structures ^29,30^.

### Csb^m/m^ and DKO mice have low acyl-CoAs which are improved by the ketogenic diet

Previously it was found that acetyl-CoA levels are reduced in CSB deficient cells^4^. We therefore decided to investigate the acetyl-CoA and related metabolites by mass spectrometry. Indeed, in the standard chow fed mice, we see a decrease in acetyl-CoA within the quadriceps of the single and double KO mice, a decrease of acetyl-CoA in the liver of Csb^m/m^ and mIndy^-/-^ mice, and increase of acetyl-CoA in the cerebellum of the mIndy^-/-^ and Csb^m/m^ single KO mice. The ketogenic diet reversed the effects for each (Figure 4J–L). Further downstream, malonyl-CoA, formed by carboxylation of acetyl-CoA and a committed step in fatty acid synthesis, is decreased DKO SD in quadriceps, decreased for all knockouts in the cerebellum, and decreased for the single knockouts in the liver (Figure 4J–L). Succinyl-CoA, a citric acid cycle intermediate that is an entry point for branched-chain amino acids (BCAA’s) valine and isoleucine, and which a depletion of leads to mitochondrial deficiency and muscle weakness^31,32^, is interestingly depleted in the mIndy^-/-^ and Csb^m/m^ SD mice, however not affected in DKO SD mice, while only WT and DKO mice are upregulated with a ketogenic diet. We also observed that glutaryl-CoA, an intermediate of lysine and tryptophan metabolism that are substrates in the synthesis of NAD, is up-regulated generally by the KD in quadriceps (Figure 4L).

### Ketones attenuate cellular features of Cockayne syndrome

We next wanted to assess molecularly whether ketones could impact cellular health outcomes and the mechanism with which this occurs. It’s known that CSB cells are also sensitive to UV damage. We observed that UV sensitive CSB cells when treated in low glucose conditions have increased UV survival (Figure 5A). Given, the UV survival improvements, we next wanted to test if cells from Cockayne syndrome patients displayed increased cellular senescence and if ketones would affect this premature aging phenotype. First, we investigated whether there was a higher incidence of senescence in primary dermal fibroblasts from a Cockayne syndrome patient or healthy control (Supplementary Figure 6A). Using a deep neural network able to predict senescence based on DAPI staining^33^ we observed that Cockayne syndrome fibroblasts are indeed predicted to be more senescent with increasing passage number than healthy control cells. Interestingly, 10mM BHB decrease senescence of control cells at low and medium passages and CS cells at the low passage (Figure 5B).

**Figure 5.**
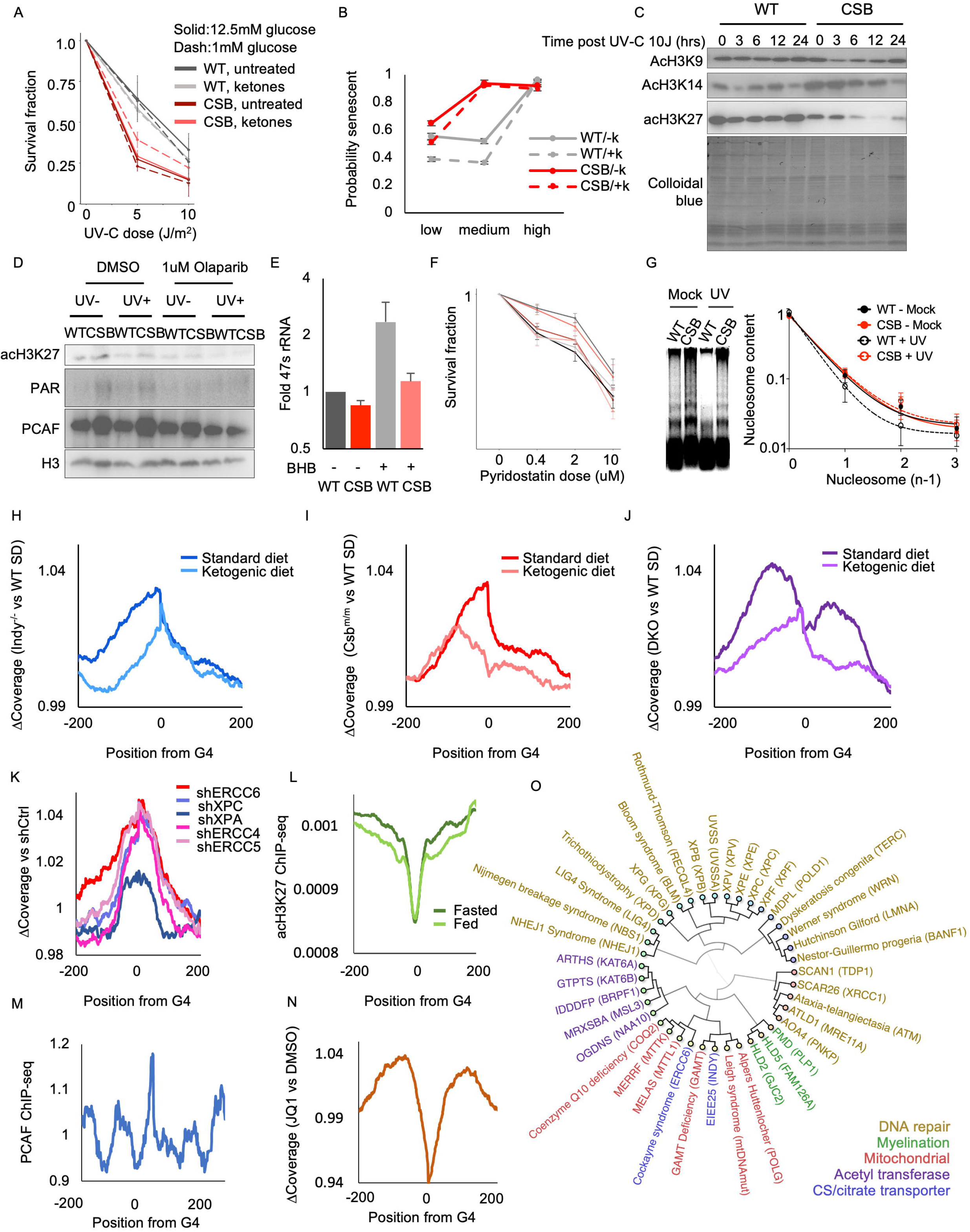
Ketones reduce transcriptional stalling at G4 structures. **A** UV-C survival of WT and CSB deficient cells (CS1AN), with low and high glucose, with and without ketones (n=3, SEM). **B** Senescence prediction of WT and CSB fibroblasts, at low (WT: p6, CSB: p4), medium (WT: p13, CSB: p13) and high passages (WT: p28, CSB: p25). **C** Western blot of histone acetylation levels in SH-SY5Y cells after exposure UV-C (10J/m2) **D** Western blot of H3 IP of acid extracted histones treated with mock (DMSO) and 1uM Olaparib (n=3, SEM). **E** 47s transcription in patient fibroblasts of WT (GM00038) and CSB (GM01428) (n=4, SEM). **F** HeLa cell survival (left) in low glucose (1mM) treated with either Pyridostatin, Pyridostatin + 1uM CX5461, or Pyridostatin + 10mM ketones; AUC of survival experiment (right) (n=5, SEM). **G** Micrococcal nuclease experiment of WT and CSB CS1AN cells, with and without UV (n=5, SEM). **H-J** Normalized coverage around all predicted G4 structures of Indy^-/-^, Csb^m/m^, and DKO mice (n=6 per group). **K** Normalized coverage around all G4 structures in cells subjected to indicated knockdowns (mined from GSE168861). **L** ChIP-seq analysis of ac-H3K27 pulldown at G4 structures (mined from GSE134044). **M** ChIP-seq analysis of PCAF pulldown at G4 structures (GSE93975). **N** Normalized coverage around all G4 structures in cells treated with JQ1 (GSE56267). **O** Hierarchical cluster of disease related to various cellular processes.

### Decreased histone acetylation in CS is recovered with ketone administration

In addition to being a substrate for mitochondrial energy metabolism, ketones also directly impact the epigenetic landscape through inhibition of histone deacetylases and as acetyl-CoA donors for acetylation reactions^34,35^. Further, it has been shown previously that RNA Pol I transcription is stimulated by acetyl-CoA^36^, a process that is deficient in Cockayne syndrome. We therefore investigated the acetylation patterns upon loss of CSB. To do this we generated CSB knockout cells using CRISPR-Cas9 targeting of exon 2 in the neurologically relevant SH-SY5Y cell lines. As expected, the knockout cells were more sensitive to UV-C radiation (Supplementary Figure 6B). We therefore investigated how DNA damage might impact histone acetylation 0, 3, 6, 12 and 24 hours after irradiation. Using this approach we observed that ac-H3K27, a marker known to regulate RNA pol I mediated transcription^37^, was depleted in CSB deficient cells (Figure 5C). Previous work has shown that the DNA damage response recruits histone acetyl transferases to the damage site ^38,39^. Notably, this histone acetyl transferase P300/CBP-associated factor (PCAF) has previously been shown to be essential for NER^38^ and involved in the metabolic adaptation of CSB deficient cells ^4^. We therefore wanted to further investigate whether decreased acetylation was linked to the DNA damage responses. Interestingly, treatment with the PARP inhibitor Olaparib decreased the levels of H3K27ac and PCAF recruitment to histones (Figure 5D). In summary, ketones attenuated UV induced cell death and induced acetylation in CSB deficient cells.

### Ketones alleviate rRNA transcription through secondary structures

Given that secondary structures more frequently occur in ribosomal DNA and these can activate the DNA damage response^29^, we investigated the role of ketones with regards to rRNA transcription. We observed that BHB increases 47S rRNA transcription significantly in dermal fibroblasts (Figure 5E), from healthy controls and CSB patients. We were curious if the impact of stabilizing G4 structures would show an additive affect with RNA Pol I inhibition. Therefore, we treated WT and CRISPR KO of CSB HeLa cells in low glucose media with the G4 stabilizer pyridostatin, in combination with the RNA Pol I inhibitor CX5461 and investigated cellular survival. Interestingly, we found non-additive effects between the treatments suggesting that both drugs elicit their toxic effect through the same mechanism (Figure 5F). We next questioned whether CSB deficient patient derived cells (CS1AN) were able to compensate under stress with chromatin decondensation, which could allow repair to take place. We observed that chromatin in CS cells were not able to decondense after UV damage (Figure 5G). Taken together, these results indicate that G4 stalling and RNA Pol I inhibition are epistatic and that chromatin condensation is increased in CSB deficient cells perhaps consistent with a general decrease in histone acetylation.

### Ketones relieve transcriptional stalling at G4 structures

Previously it was shown that stalling of transcription occurs at various types of secondary DNA structures particularly in CSB deficient cells, with the strongest stalling occurring where G-quadruplex structures are present^29^. Strikingly, by analyzing our RNA seq data we discovered stalling of transcription at G-quadruplexes in mIndy^-/-^ (Figure 5H) and Csb^m/m^ (Figure 5I) and was particularly prominent in DKO and this stalling was normalized by a ketogenic diet (Figure 5J, Supplementary Figure 7). To further explore these effects on transcription we datamined several studies. First, we explored if inhibiting various stages of nucleotide excision repair by knocking down ERCC6, XPC, XPA, ERCC4, and ERCC5 would impact stalling at G4 structures (GSE168861). Notably, we observed each of the knockdown cell lines experience increased stalling around the G4 sites with the strongest stalling occurring after CSB knockdown (Figure 5K). Since transcription is heavily affected by histone acetylation, we speculated that ketones might impact stalling at these structures. Indeed, in a published H3K27 acetylation ChIP-seq data (GSE134044) we found that H3K27 acetylation is lost at G4 structures and that fasting, an intervention leading to ketosis, normalized H3K27 acetylation at predicted G4 structures. (Figure 5L). Further, in a published ChIP-seq dataset (GSE93975) we found PCAF enrichment at G4 structures (Figure 5M). And, by inhibiting transcriptional elongation with JQ1, a BET bromodomain inhibitor, (GSE56267) leads to increase stalling at G4s (Figure 5N). To understand if this has clinical relevance, we identified diseases with defects in acetylation, myelination and mitochondrial function and clustered their clinical phenotype as previously described ^29,40^. Interestingly, we found close phenotypical overlap with Cockayne syndrome and Early infantile encephalopathy and epilepsy 25 (EIEE25) caused by mutations in INDY in all these diseases (Figure 5O). These data indicate that ketones may increase histone acetylation leading to transcriptional readthrough of secondary DNA structures and a reduction in PARP activation (Supplementary Figure 8).

## Discussion

Here, we found that manipulating the acetyl-CoA donors impacts age-related phenotypes in models of Cockayne syndrome. Whether drosophila contained key NER proteins such as CSB and CSA was debated and doubted in the past^41^. However, recent evidence suggests that TC-NER does take place in S2 drosophila cell lines and a homologue of the XPA binding protein 2 (XAB2) has recently been found in drosophila^42–44^. With Hel89B as the potential homolog to ERCC6, this finally provides a mechanistic basis for TC-NER in this species. Interestingly, Hel89B was also shown to have homology to BTAF1^45^, which can act as a transcriptional repressor as well. Nevertheless, Hel89B deficiency appear to recapitulate two key hallmarks of TC-NER deficiency: UV-sensitivity and decreased resumption of RNA synthesis after UV-C irradiation. Furthermore, flies with less Hel89B live significantly shorter than their wild-type counterparts. These are all classical Cockayne syndrome phenotypes and provide evidence for a potentially novel ERCC6 homologue and the usability of Hel89B deficient flies as a new Cockayne syndrome model.

In mammals, loss of mIndy in Csb mice exacerbates the phenotype at the behavioral, metabolic and transcriptomic level and ketones rescue these phenotypes. Interestingly, it was recently reported that DNA damage increases β-oxidation as a means of producing acetyl-CoA ^11^. This would be consistent with our findings here and previously ^4^ where loss of Csb leads to increased fatty acid oxidation. We hypothesize, that the switch to fatty acid oxidation is an adaptive response to allow increased acetylation to take place in response to DNA damage perhaps to facilitate histone relaxation and increased DNA accessibility.

The core molecular mechanism with which ketones confer neuroprotection and longevity has not yet been resolved. In Cockayne syndrome models, it was suggested that the effectiveness of ketones could be mediated trough activation of SIRT1^4^. Ketones are also shown to increase histone acetylation and act as an HDAC inhibitor^35^. We further validated these findings with ketones increasing overall histone acetylation levels in both our cell and mouse models of Cockayne syndrome. At the molecular level, it appears that ketones are able to facilitate transcription through G4 structures by increasing histone acetylation. This is particularly prominent at rRNA where H3K27 acetylation and acetyl-CoA levels are known to positively regulate transcription. Thus, ketones likely facilitate histone acetylation thereby allowing chromatin decondensation and access for transcription associated helicases that can unwind G4 structure. Notably, loss of transcription associated helicases XPD and XPB, two helicases known to bind G4 structures, can also cause Cockayne syndrome ^46^. At a more general level, our data provide evidence that ketones could be neuroprotective by allowing resolution of secondary DNA structures.

## Materials and Methods

### Animals and diets

Six-month-old-mice were fed a standard AIN-93G diet ad libitum or a ketogenic diet (Dyets) with 60% calories coming from fat, primarily hydrogenated coconut oil. WT, mIndy^-/-^, Csb^m/m^, and mIndy^-/-^/ Csb^m/m^ were of C57BL6 background and generated by first mating single knockouts to generate double heterozygotes. Double heterozygotes were then mated to generated all genotypes that occurred at roughly mendelian frequencies. Unless otherwise stated measurements took place at 6 months, 9 months and 12 months of age. Body weight, chipped body temperature via electronical tagging (Biomedic Data System, Maywood, NJ), and food consumption were recorded every two weeks of the study after the diet was administered. Animal rooms were maintained at 20-22°C and a 12-hour light/dark cycle. All animal protocols were approved by the Animal Care and Use Committee (277-TGB-2019) of the National Institute on Aging (Baltimore, MD, USA).

### Metabolic assessment

The metabolic rate of the mice was assessed by indirect calorimetry in open-circuit oxymax chambers using the Comprehensive Lab Animal Monitoring System (CLAMS; Columbus Instruments, Columbus). Mice were singly housed with water and food available ad libitum and maintained at 24°C under a 12:12 h light-dark cycle (light period 0600-1800).

### Body Composition

Measurements of lean, fat and fluid mass in live mice were acquired by nuclear magnetic resonance (NMR) using the Minispec LF90 (Bruker Optics, Billerica, MA).

### Gait analysis

Gait analysis was performed previously as described^27^ (PMID 30649206). In brief, mice were acclimated a week before in the TSE MotoRater, in order to assure smooth running across the narrow platform. When ready for the experiment, mice were recorded running over the path twice, and this data was averaged for analysis.

### Startle Chamber analysis

The startle response system (SR-Labs, San Diego, California, USA) was used to assess hearing in the mice. Mice were acclimated before the sound cycle began.

The cycle consisted of 20ms pulses of 65, 70, 75, 80, 85, 90, 95, 100, 105, 110, 115, and 120 dB, with an inter-trial interval (ITI) of 100 ms.

### Oral Glucose Tolerance Test (OGTT) and Insulin Tolerance Test (ITT)

Mice were fasted for 3 hours prior to OGTT and ITT. For OGTT, mice received a 30% glucose solution (1.5g/kg glucose by gavage). For ITT, a dose of 1IU/kg insulin (Novo Nordisk) was injected intraperitoneally. Blood glucose was measured at time points 0, 15, 30, 60, and 120 minutes following gavage or injection.

### Metabolic treadmills

Mice were acclimated on the metabolic treadmills (Columbus Instruments) at 5 m/min for 30 min the day prior to testing to ensure familiarity. The next day, mice warmed up on the treadmill with 5 m/min for 2 minutes, and then ran at 12 m/min for 10 minutes. VO_2_ and VCO_2_ were simultaneously recorded.

### Histology

Histology and mounting of the liver and cerebellum was performed by the Core Facility for Integrated Microscopy (CFIM) with formalin fixation followed by paraffin embedding and section. The slides were imaged in CFIM as well, using 20x magnification in the automated slide scanner Axio Scan.Z1 (Zeiss). Liver was stained with H&E and cerebellum was stained with either H&E or luxol fast blue.

### RNA-sequencing and RNA extraction

RNA isolation was performed using TRIzol (Thermo-Fischer). RNA-seq was performed on 48 mouse cerebellum samples, by BGI genomics using whole RNA seq analysis, rRNA depleted. Analysis was performed using an in-house pipeline described:

Paired-end reads from 44 RNA-Seq samples were aligned to mm9 using bowtie2^47^. Differential expression analysis was performed using a Salmon^48^ → tximeta^49^ → DESeq2^48^. GSEA was performed directly on DESeq2 normalized counts. All comparisons were made against WTSD. The Gene Set used was the C5 collection based Gene Ontology (GO) terms^50,51^. Terms were filtered for FDR < 0.05 and sorted by Normalized Enrichment Score (NES).

Paired-end reads from 44 RNA-Seq samples were aligned to mm9 using bowtie2^47^. We used a list of identified regions from the non-B DB^52^ database predicted to form G-quadruplex candidate structures to delineate a 1000bp window (500bp before start of motif and 500bp after end of motif, not including motif itself.) We then calculated the allelic depth surrounding G-quadruplex motifs within the 1000bp window. We specifically selected G-quadruplex motifs for which the entire 1000bp is contained within gene sequences. Since G-quadruplex motifs are stranded we distinguish sense transcription from antisense transcription. We then calculated the reference allelic depth and the reference allele frequency. Code can be found at github.com/scheibye-knudsen-lab/INDYCSB-mutations.

### Metabolomics quantification and analysis

Metabolomics measurements were performed on 48 mouse cerebellum samples by Scripps Center for Metabolomics (La Jolla, California, USA) using untargeted metabolomics, HILIC negative. The analysis was performed using a combination of the XCMS platform and MetaboAnalystR 2.0.

### Acyl-CoA extraction and LC-MS of mice tissue

Analysts blinded to experimental groups performed acyl-CoA analysis by stable isotope dilution liquid chromatography-high resolution mass spectrometry on an Ultimate 3000 autosampler coupled to a Thermo Q-Exactive Plus as previously described^53^. In brief, frozen samples and calibration curve points were spiked with 100 μL of [^13^C_3_^15^N_1_]-acyl-CoA internal standards generated as previously described from [^13^C_3_^15^N_1_]-pantothenic acid^54^, samples were homogenized by probe tip sonication, then acyl-CoA extraction was performed by solid phase extraction^55^.

### Drosophila strains

The following Drosophila strains are described in the following references: UAS-mGFP, UAS-mCherry, yellow-white (yw), and Actin-Gal4/CyO were kindly provided by Hector Herranz’s group at ICMM. The following stocks were provided by the Bloomington *Drosophila* Stock Center: Hel89B #1 (#32895), w1118 (#5905), Indy^206^ (#27901), and Indy^302^ (#27902). The following stocks were provided by the Vienna Drosophila RNAi Center: Hel89B #2 (#4237). The w;Actin-Gal4/CyO;Indy^206^/Bal and w;Actin-Gal4/CyO;Indy^302^/Bal driver lines were generated in-house. Flies were maintained on standard SYA food at 25C in a 12 hour light/dark cycle with constant 60% humidity. Ketones used for all experiments were (R)-3-Hydroxybutyric acid (Sigma-Aldrich #54920). Citrate used for all experiments was sodium citrate (Sigma-Aldrich #1613859). Olaparib used for all experiments was Selleckchem #AZD2281.

### UV survival assay

Third instar larvae were exposed to UV-C irradiation (custom built machine) at log-scale doses: 0J/m^2^, 50J/m^2^, 100J/m^2^, and 200J/m^2^. Afterwards, they were placed in standard agar food in standard vials, and survival was recorded in subsequent days. Flies were also video recorded throughout their hatching.

### Lifespan and motor function throughout the life

Drosophila crosses laid eggs in 25C and adult experimental flies were allowed to emerge and mate for 1 day. Afterwards adult flies were lightly anaesthetized with CO_2_ and males selected to assess for experiments. Per vial, a density of 10 flies were used for lifespan assessment. Fruit flies were recorded every hour of every day using the platform from Tracked.bio. Each week, manual counts were also taken of the flies. Flies were fed with sufficient amounts of food for 1 week and therefore fresh food was exchanged weekly. Food contained low melting agar (OmniPur® Agarose, Low Melting, CAS 9012-36-6, Merck) during lifespan assessments only to ensure the quality of compounds was preserved. Lifetime motor function was calculated as the the area under the curve of the linear regression of max distance traveled per day; this larger time point used to calculated AUC was the mean lifespan of the flies.

### Resumption of RNA synthesis

S2 cells were UV’ed with 40J/m^2^ of UV-C (custom built machine) and then incubated in 1 mM EU in the culture media for 3 hours followed by processing with the Click-IT RNA Alexa Fluor 594 imaging kit (ThermoFisher #C10330).

### RT-PCR of third instar larvae

3^rd^ third instar larvae were selected after being grown in drosophila fly cages with media and yeast paste. The larvae were RNA extracted with a standard TRIzol, phenol:chloroform protocol. Before extraction, flies were pulverized with 3D printed spears. cDNA synthesis was performed with the Applied Biosystems™ High-Capacity cDNA Reverse Transcription Kit (#4368814). RT-PCR was performed with SensiFAST™ master mix. Primer sequences used were: Hel89B_forward: CCGCCTTGAAGCAACTTCTC, Hel89B_reverse: CCTCGTACTGATGCTCGGAC, rp49_forward: AAGCGGCGACGCACTCTGTT, and rp49_reverse: GCCCAGCATACAGGCCCAAG.

### RT-PCR of S2 cells

S2 cells were cultured in 35mm dishes and treated with the respective RNAi or scramble RNAi. RNA was then extracted with the Zymo Research RNA MiniPrep Plus kit (#R2072). cDNA synthesis was performed with the Applied Biosystems™ High-Capacity cDNA Reverse Transcription Kit (#4368814). RT-PCR was performed with SensiFAST™ master mix. Oligo sequences used for knockdown experiments were: siHel89B1: AAGAUCCUUACUCUAGAUCAA, siHel89B2a: AAUGAAGGAUCUGCAGGCUA, siHel89B2b: CACCUCACAGAUCUUUGACC. siHel89B1 showed the greatest knockdown efficiency and was used for further experimentation.

### Acyl-CoA extraction and LC-MS of fruit flies

Female fruit flies were collected at the indicated time points. Analysts blinded to experimental groups performed acyl-CoA analysis by stable isotope dilution liquid chromatography-high resolution mass spectrometry on an Ultimate 3000 autosampler coupled to a Thermo Q-Exactive Plus as previously described^53^. In brief, frozen samples and calibration curve points were spiked with 100 μL of [^13^C_3_^15^N_1_]-acyl-CoA internal standards generated as previously described from [^13^C_3_^15^N_1_]-pantothenic acid^54^, samples were homogenized by probe tip sonication, then acyl-CoA extraction was performed by solid phase extraction^55^.

### Negative geotaxis assay

Fruit flies were collected at 5 days of age. A custom 3D printed apparatus was fabricated to perform this experiment. Two drops were used to fully send the flies to the bottom of the vials and the flies were recorded while they ascended to the top of the vials. The flies were tracked and quantified with custom software in Python 3.

### Hatch rate

Hatch rate was measured by crossing all the fly genotypes at 15 males and 9 females ratios. The flies were then separated by gender and genotype based on balancers, and percentage hatched was quantified based on the desired male genotype/phenotype for the lifespan experiments.

### Cell culture

HeLa and SH-SY5Y CSB CRISPR-Cas9 knockouts were generated with the following plasmid: the plasmid used was modified from a pCas-Guide-EF1a-Cherry from Origene, modified to contain a guide RNA targeting the PARP binding domain of CSB, sequence: GATCGCATCGACCGACATCAGATCCG. In brief, the cells were generated by initial sorting of Cherry-positive cells by flow cytometry and single clones were isolated by serial dilution in a 96-well plate. The HeLa cells were cultured in Dulbecco’s Modified Eagle Medium (DMEM), High glucose, GlutaMAX™. The SH-SY5Y cells were culture in a mixture of half Hams F12 media (Biowest # L0136) with 10% FBS and 1% pen-strep, and half DMEM high glucose, GlutaMAX™ Supplement, pyruvate (Fischer #12077549). WT (GM00038) and CSB (GM01428, GM10903) fibroblast cell lines were obtained from the Coriell Institute (Camden, New Jersey, USA), and cultured in MEM (with Earle’s salts, without L-glutamine, with NEAA; VWR # L0430-500) media with 15% FBS and 1% pen-strep. Unless otherwise stated, high glucose media for experiments was 25mM and low glucose media was 1mM. Low glucose media was prepared by mixing 25mM media with no glucose DMEM (Fisher #11520416) to the correct proportions.

### RT-PCR of 47S

Cells were seeded at a density of 60,000 cells per well in 6-well plates. The cells were RNA extracted with a standard TRIzol, phenol:chloroform protocol. RT-PCR was performed with SensiFAST™ master mix. 47S was probed with the following primer sequences: 47S_forward: CGGGTTATTGCTGACACGC and 47S_reverse: CAACCTCTCCAGCGACAGG, which were adapted from a previous study (PMID: 31841110). The GAPDH sequences used were: GAPDH_forward: GTCAGCCGCATCTTCTTTTG and GAPDH_reverse: GCGCCCAATACGACCAAATC.

### UV survival

CS1AN WT and CSB cells were plated at a density of 30,000 in 24-well plates and pre-treated with or without ketones and and with or without low glucose media for 24 hours. After 24 hours, the cells were UV’ed at the indicated doses.

### UV histone acetylation

SH-SY5Y WT and CSB cells were plated at density of 3,000,000 cells per dish. 24 hours later they were UV irradiated with 10J/m^2^ and then collected at time points of 0, 3, 6, 12, and 24 hours. Cells were then acid histone extracted (PMID: 17545981)

### Immunoprecipitation of H3

Samples were added to IP buffer (150mM NaCl, 1% NP-40, 2mM EDTA, 25mM Tris pH 8.0) supplemented with protease inihibitor, phosphatase inhibitor and sodium butyrate. 10ug of H3 antibody was then added, and incubated for 1 hour on ice. Then 70uL a/g beads (Santa Cruz Biotechnology sc-2003) were added and tumbled overnight at 4C. The next day, the samples were washed 3 times with 500uL of IP buffer: they were spun down each time at 400g for 5 minutes at 4C. After washing, 100uL of 1x Laemmli buffer was added to the beads.

### Micrococcal Nuclease

The protocol to perform the micrococcal nuclease was adapted (PMID: 23396441). In brief, CS1AN cells were trypsinized wash in ice cold PBS containing 5mM sodium butyrate and then centrifuged at 600g for 3 minutes. The cells were lysed in hypotonic buffer (10mM Tris–HCl pH 7.4, 10mM KCl, 15mM MgCl_2_, 5mM Na butyrate) on ice for 10 min. The pellet’s nuclei was centrifuged at 1000g for 5 minutes. The nuclei were then resuspended in Micrococcal nuclease (MNase) digestion buffer [0.32 M sucrose, 50 mM Tris–HCl (pH 7.5), 4mM MgCl_2_, 1 mM CaCl_25_, 0.1mM phenylmethylsulfonyl fluoride (PMSF), 5mM sodium butyrate]. MNase was then added and incubated at 37°C. The MNase reaction was stopped with 10mM EDTA. The pellet was then resuspended in MNase digestion buffer supplemented with 10mM EDTA and RNase and incubate at 37°C for 30 minutes. STOP buffer (proteinase K, SDS, EDTA) was then added at 37°C for 30 minutes. The DNA was finally extracted and purified adapted from the Thermo Fisher Ethanol DNA Purification protocol. In brief, glycogen (20ug/uL), 7.5M NH_4_OAc, and 100% ethanol were added to the sample in that order. The samples were then placed in −20C overnight to allowed precipitation. The next day, the sample was centrifuged at 4C for 30 minutes at 16,000g. The supernatant was then removed and combined with 150uL 70% ethanol. Next it was centrifuged at 4C for 2 minutes at 16,000g. The supernatant was again centrifuged at 4C for 30 minutes at 16,000g. The pellet was then dried and resuspended in water. The samples were then analyzed on a 1.4% agarose gel in SYBR gold.

### Population doubling

Fibroblasts were cultured in the respective treatments: untreated, 10mM ketones, and 1mM citrate. Each week the cells were passaged, counted and replated. Population doubling (PD) was calculated with the formula: PD = (Previous week PD) + 3.332*(log(cells in dish) – log(cells seeded)).

### Beta-gal measurement and quantification

Cells were stained using a beta-gal kit (Sigma-Aldrich #CS0030) and were imaged using high-content microscopy (IN Cell Analyzer 2000) with a 20x objective and INCell investigator software.

### Antibodies and reagents

Antibodies included acH3K9 (Cell signaling #9649), acH3K14 (Sigma-Aldrich #07-353), acH3K27 (Cell signaling #8173), H3 (Abcam #ab1791), PAR (Trevigen #4335-MC-100), PCAF (Cell signaling #3378), CSB (Santa Cruz Biotechnologies # sc-398022), and GAPDH (Origene #5G4-6C5). Secondary antibodies include Anti-rabbit IgG, HRP-linked antibody (Cell signaling #7074), and Anti-mouse IgG (Sigma-Aldrich #A4416). Staining for colloidal blue was from Invitrogen (Invitrogen # LC6025).

### Statistical analysis

All statistics were performed in R and GraphPad Prism 9.0. Two-way ANOVA was performed on the mouse categorical data.

## Author’s Contributions

M.A.P. Wrote the article, performed experiments, analyzed data, developed methodology; L.M.C.M, performed experiments; T.T. performed experiments; S.K. performed experiments; S.R. performed experiments; D.B. performed experiments; G.K. performed experiments; B.O. performed experiments; S.J.M. performed experiments; S.H. performed experiments; J.K. performed experiments; I.A. performed experiments; A.A.T. performed experiments; I.H. developed methodology and analyzed data; J.M. analyzed data; M.B.E. analyzed data; G.M. performed experiments; E.V. performed experiments; B.F. performed experiments; EvK performed experiments; N.W.S. supervised CoA experiments; H.H. supervised drosophila experiments; RdC supervised mouse experiments; M.S.K. Conceived the idea of the study, supervised experiments.

## Acknowledgements and Funding

MSK supported by the Novo Nordisk Foundation Challenge Programme (#NNF17OC0027812), the Nordea Foundation (#02-2017-1749), the Neye Foundation, the Lundbeck Foundation (#R324-2019-1492), the Ministry of Higher Education and Science (#0238-00003B), VitaDAO and Insilico Medicine. NWS (R01GM132261 and R01CA259111).

## Conflicts of Interest

All authors do not have conflict of interest to declare.

## Supplementary Figures

**Supplementary Figure 1**. **Drosophila phenotyping. A** Hatch rate of the genotypes used. **B** Schematic of UV-survival assay. **C** Hel89B gene expression of UAS-mGFP (control), Hel89B RNAi #1, and Hel89B RNAi #2 third-instar larvae. **D** Hel89B gene expression of siSCR (control), siHel89B #1, siHel89B #2a, and siHel89B #2b in drosophila S2 cells. **E** Acyl-CoA species of Hel89B and mGFP flies at various ages.

**Supplementary Figure 2. Lifespan curves of genotypes.** Lifespan curves of all groups depicted in one graph, n = 3 sets, 5644 flies.

**Supplementary Figure 3. Additional mouse data. A** Schematic illustration of the mouse experimental procedure. Six-month WT, mIndy^-/-^, Csb^m/m^, and DKO mice were fed either a standard or ketogenic diet for six months (12 per group), and assessed at three time points (beginning, middle and end) for behavior and metabolic readouts. At the last time point, the mice were sacrificed, tissues harvested and saved for post-mortem analysis. **B** Weights of organs. **C** Haematology analysis. **D** Open field assessment for 30 minute periods. **E** Y-maze for 10 minute periods. Spontaneous alteration tests. **F** Elevated zero maze for 5 minute periods. **G** Gait analysis using the TSE MotoRater. n=12 per group, SEM.

**Supplementary Figure 4. Omics data from the murine experiments. A** Principal component analyses of gene expression in the cerebellum by RNA seq (n=6 per group). **B** Venn diagram of altered GO terms from the RNA seq experiment. **C** Hierarchical clustering of z-score normalized metabolites in the cerebellum. **D** Top 50 and bottom 50 metabolites in heatmap.

**Supplementary Figure 5. Altered metabolic pathways in the cerebellum of mice.** n= 6 per group. All data was log-normalized and the groups are compared to the WTSD mice.

**Supplementary Figure 6. Data from csb deficient cell lines. A** Population doubling of human fibroblasts: GM00038 (WT, control), GM01428 (Cockayne syndrome), and GM10903 (De Sanctis-Cacchione syndrome). **B** Western blot confirming CSB deletion in both HeLa and SH-SY5Y cells (top), as well as UV-dose response curves (bottom). **C** Morphometric data from the senescence prediction analysis of WT and CSB fibroblasts.

**Supplementary Figure 7. Multiple DNA structures elicit transcriptional stalling in prematurely aged mice.** Coverage surrounding Non-B DNA structures of WT, Csb^m/m^, mIndy^-/-^, and DKO mice from RNA-seq. STR: Short tandem repeat, IR: inversed repeat, APR: A-phased repeat, MR: Mirror repeat, DR: Direct repeat.

**Supplementary Figure 8. Putative model of how ketones might facilitate transcriptional readthrough of secondary DNA structures.**

## Notes

### Competing Interest Statement

The authors have declared no competing interest.

