## Supplementary figures and images for "Ketones facilitate transcriptional resolution of secondary DNA structures in premature aging"

### Supplementary figures

#### Supplemental Figures

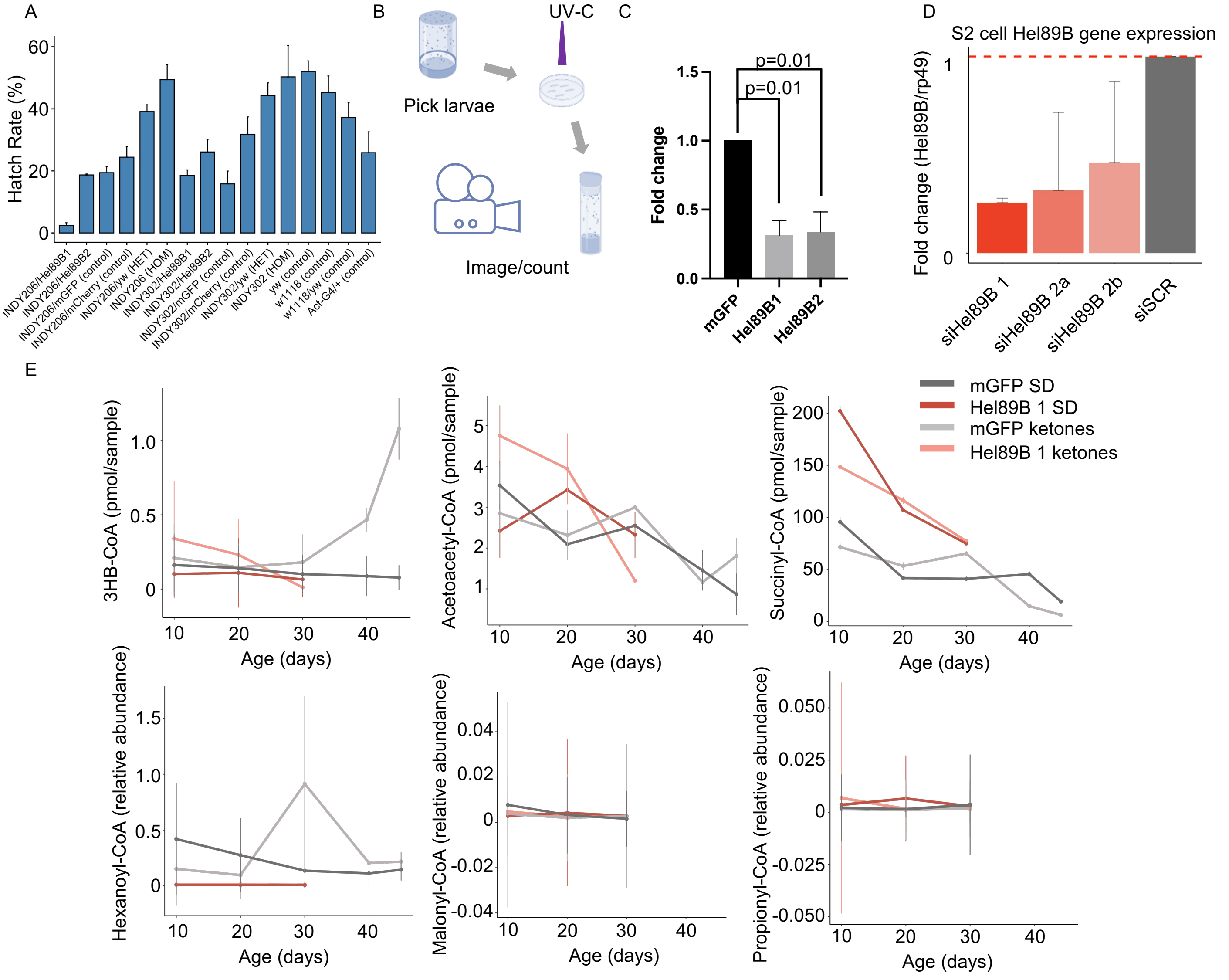

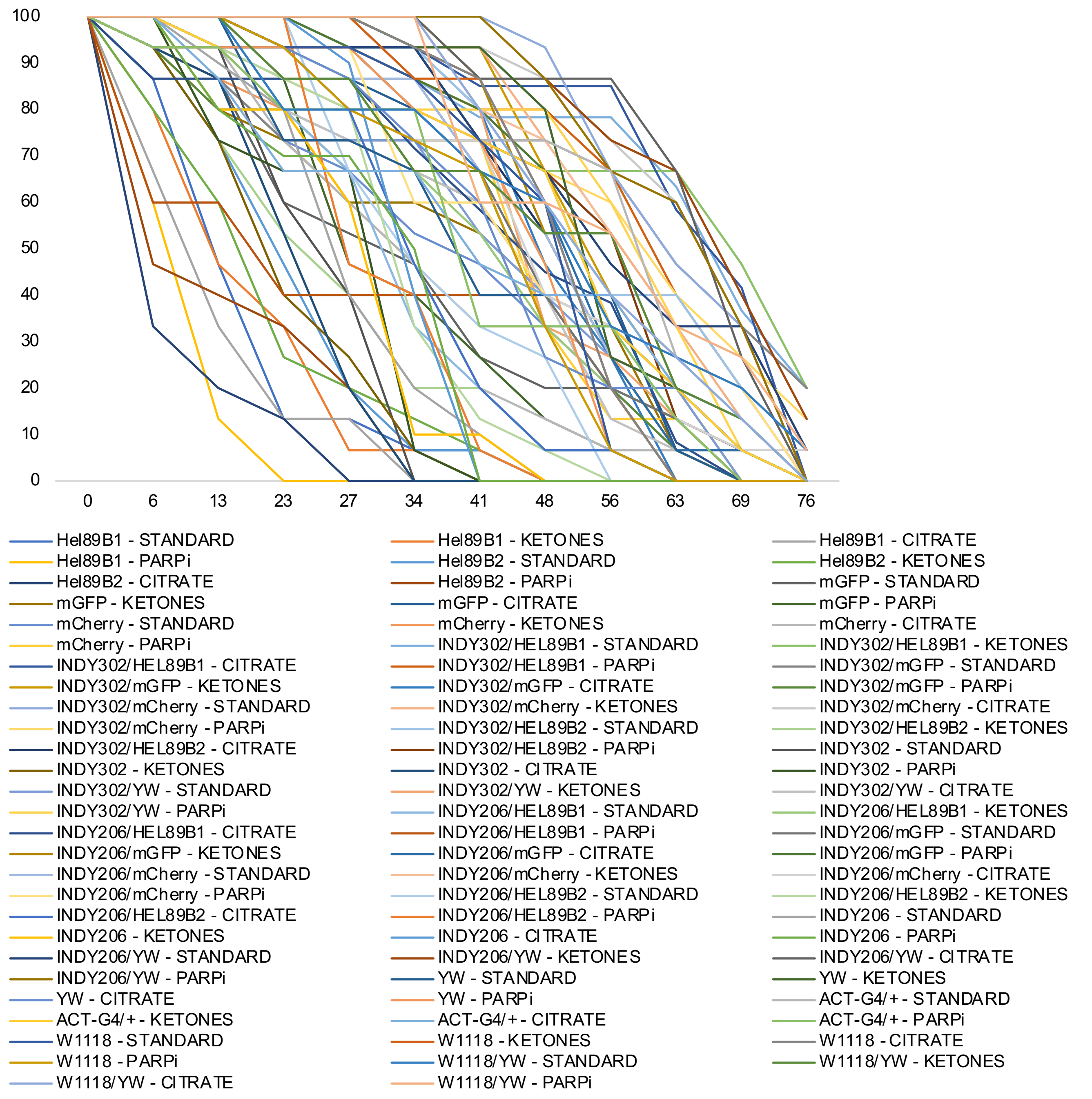

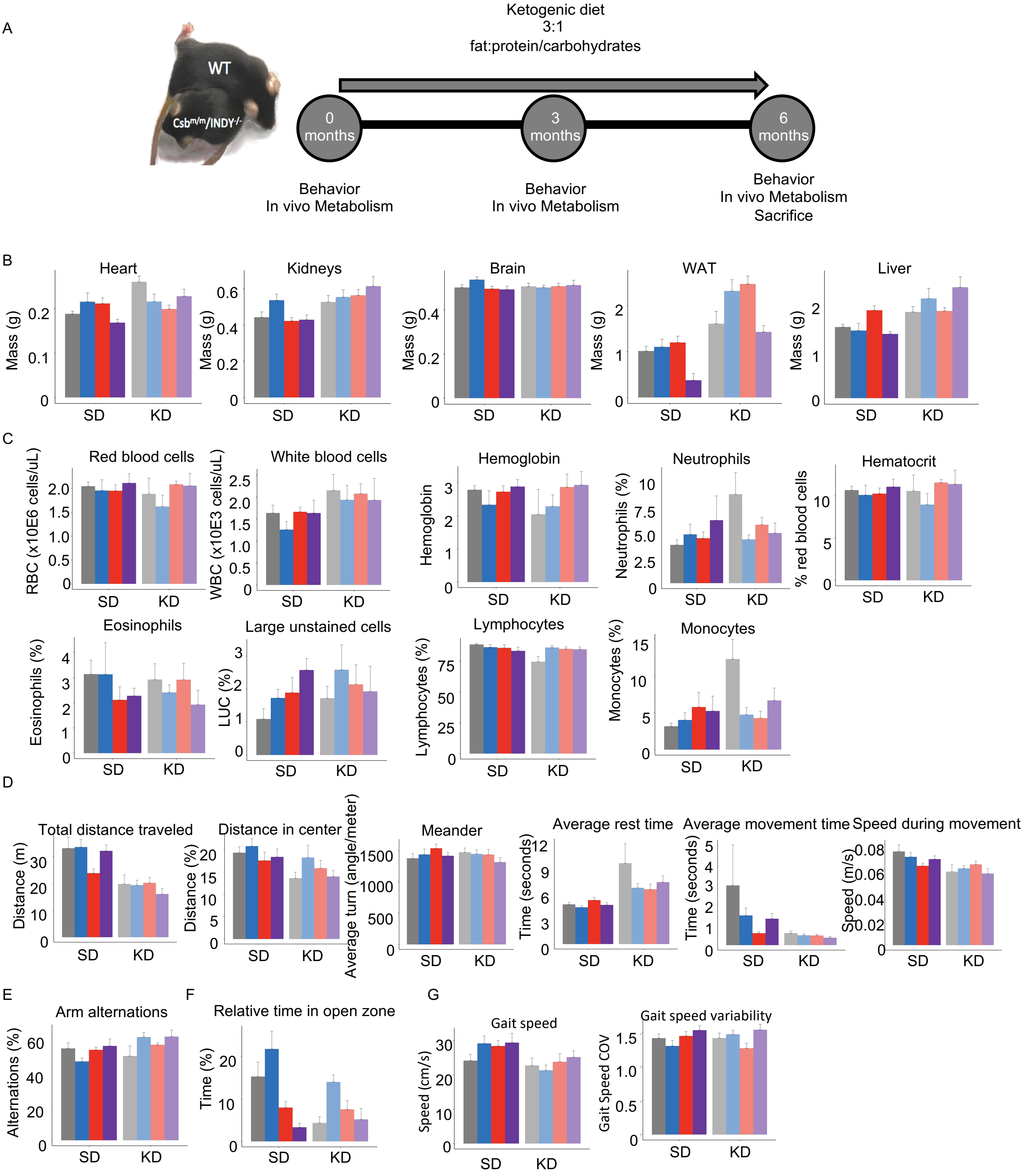

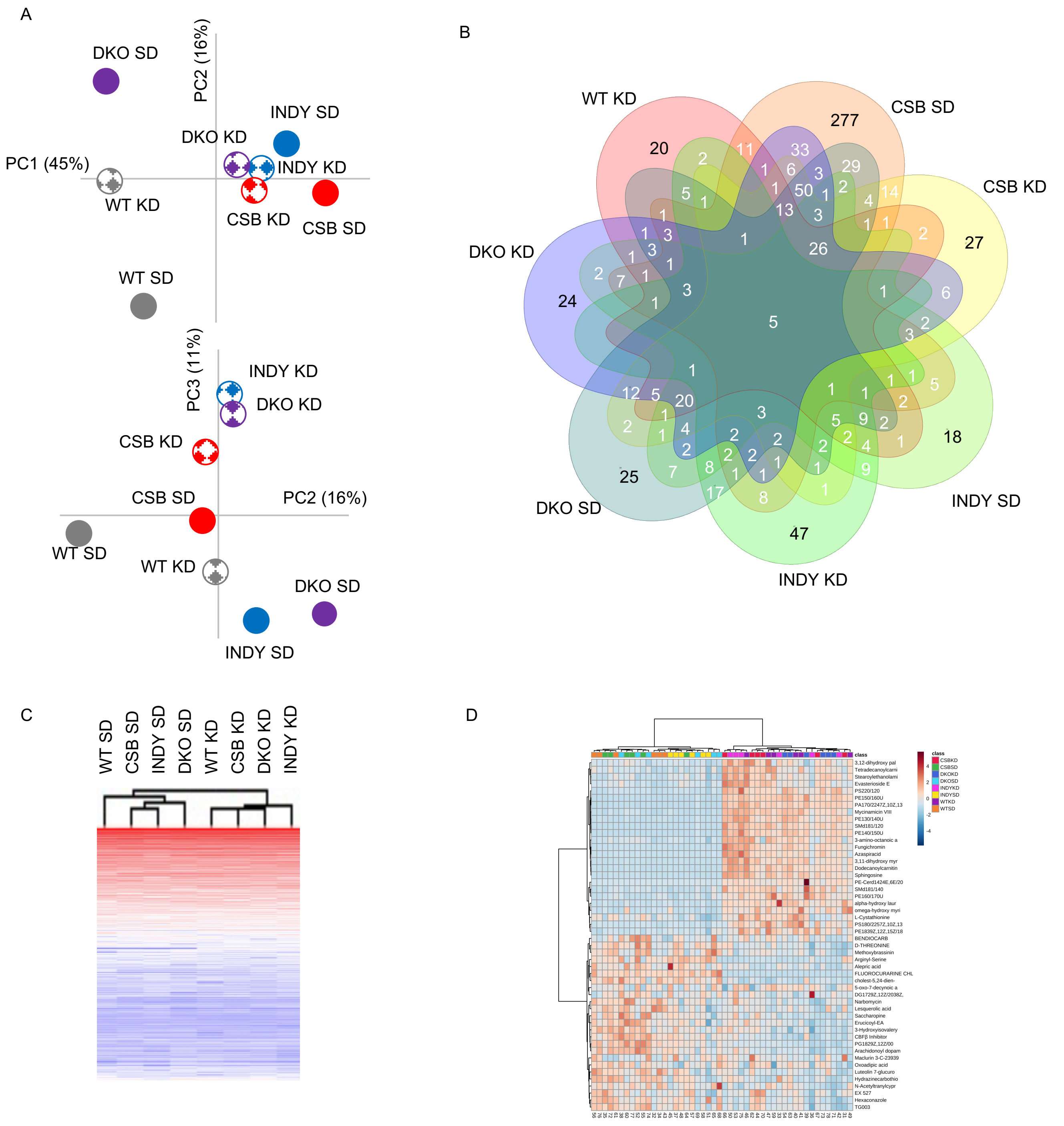

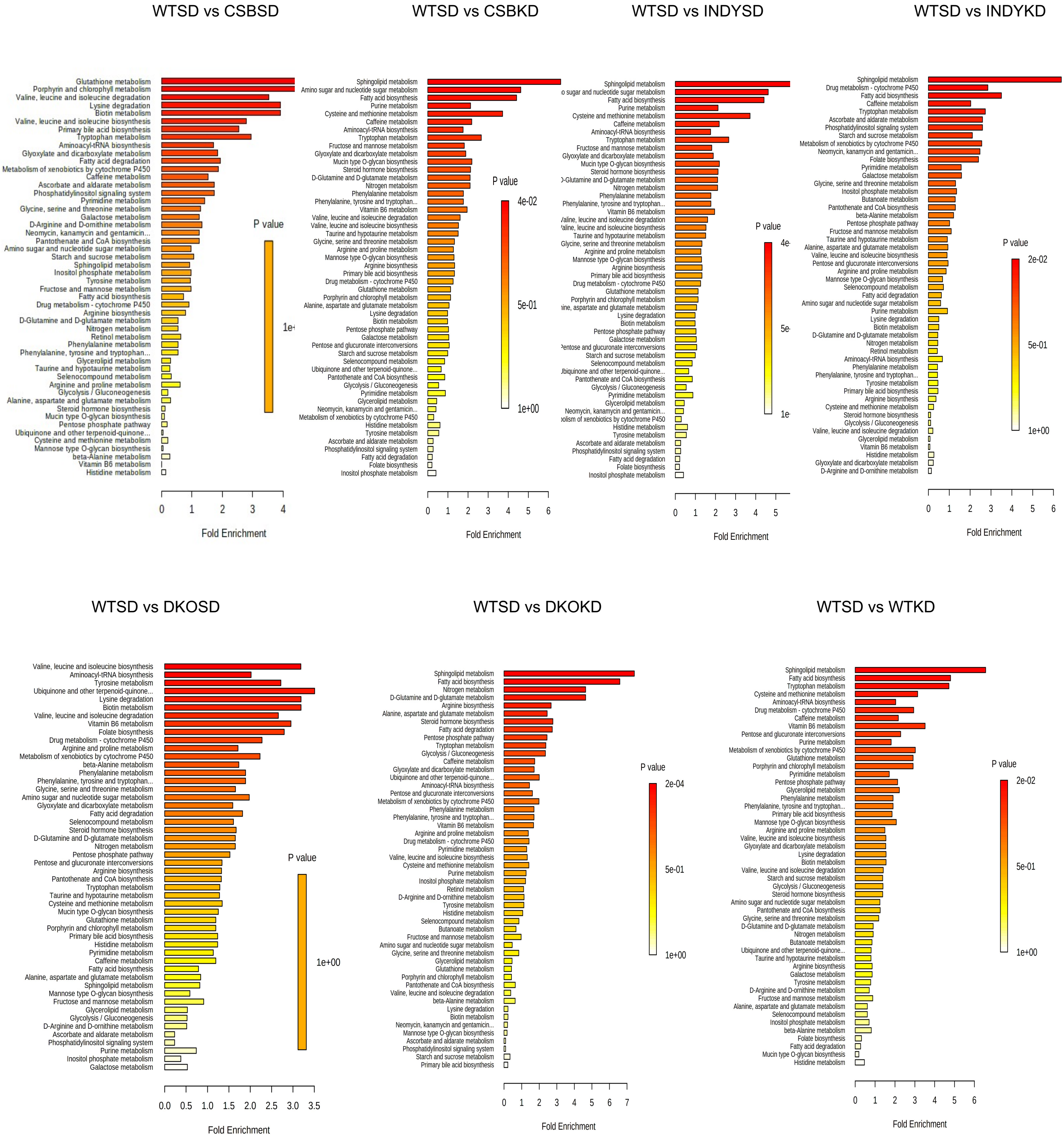

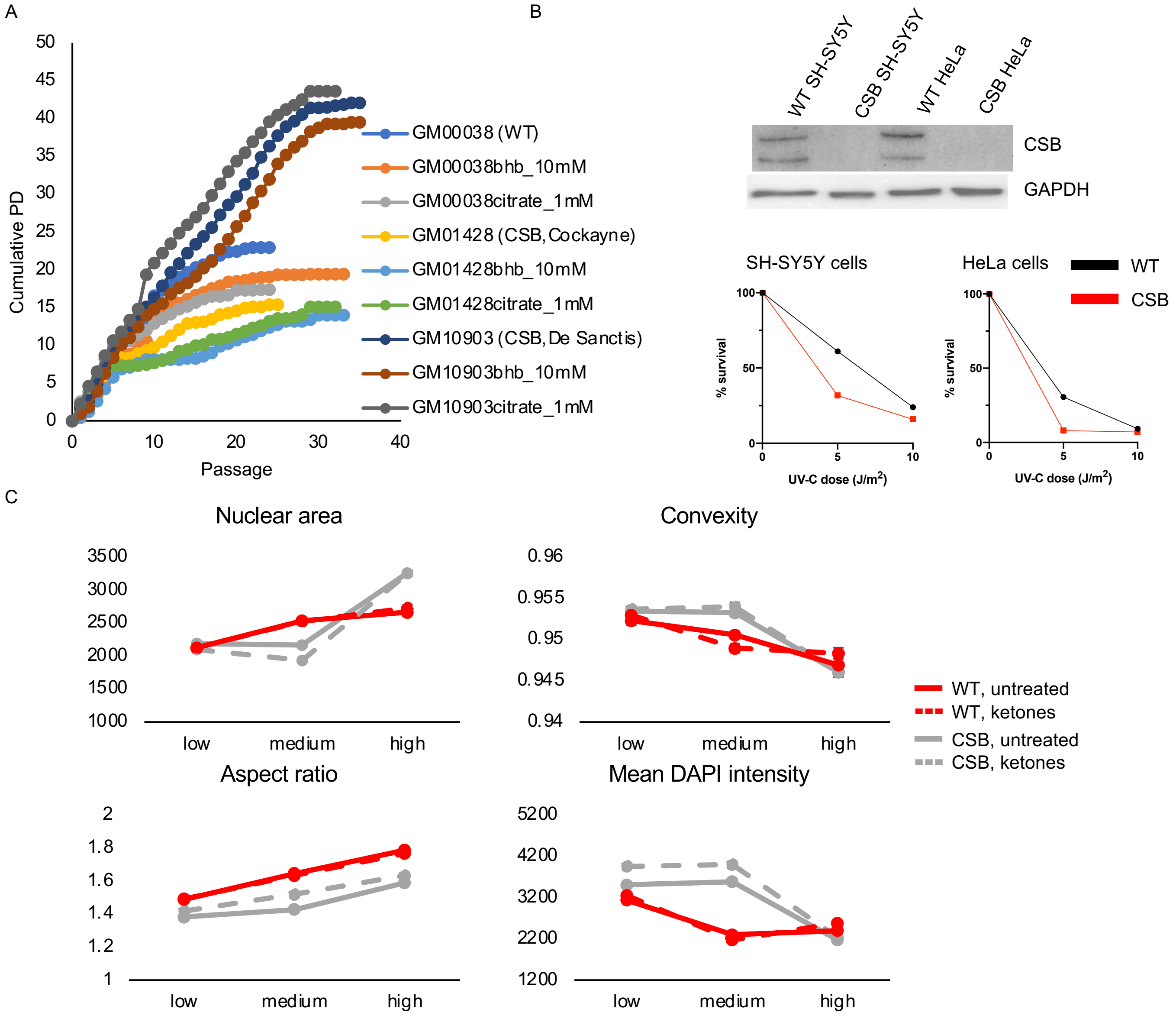

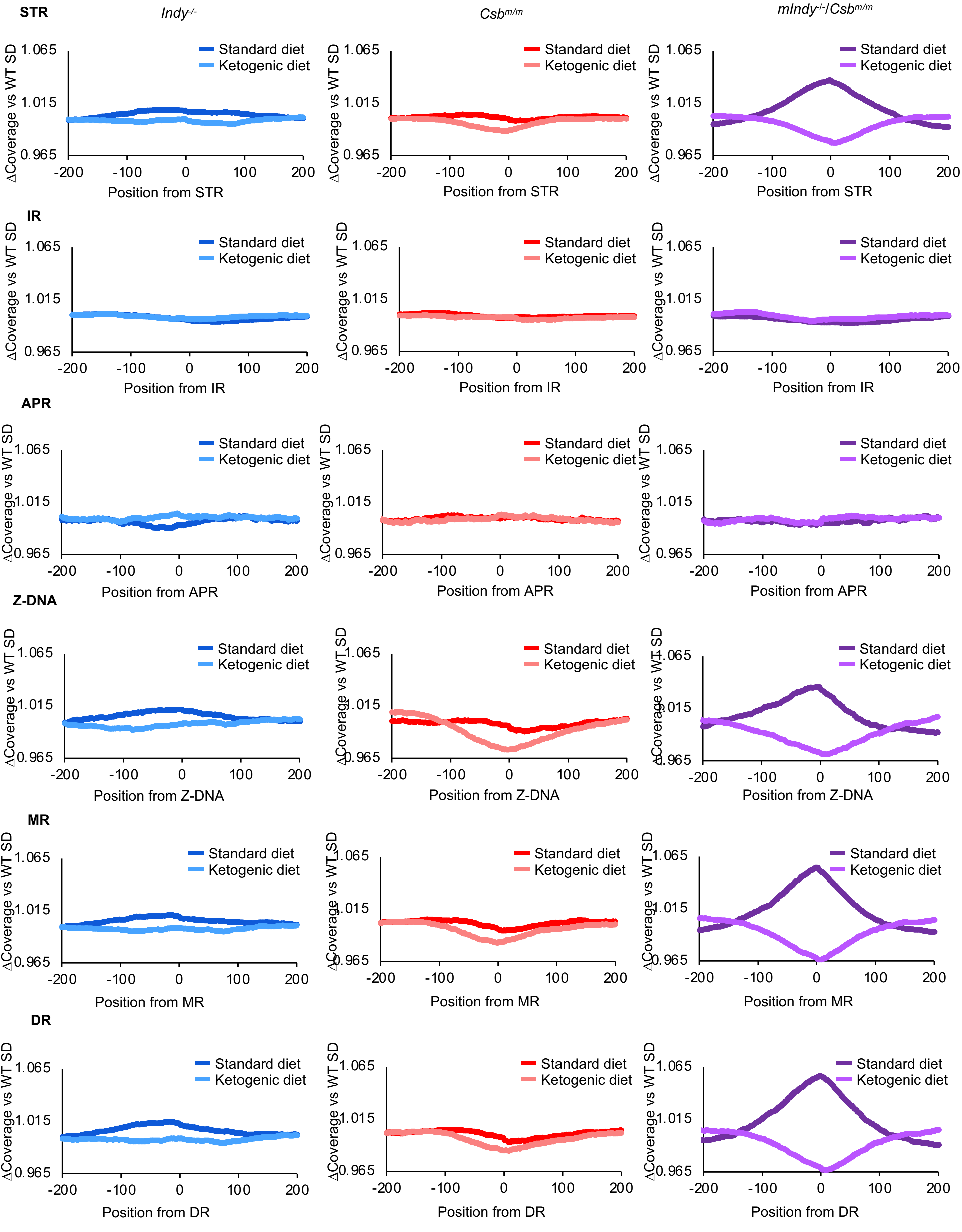

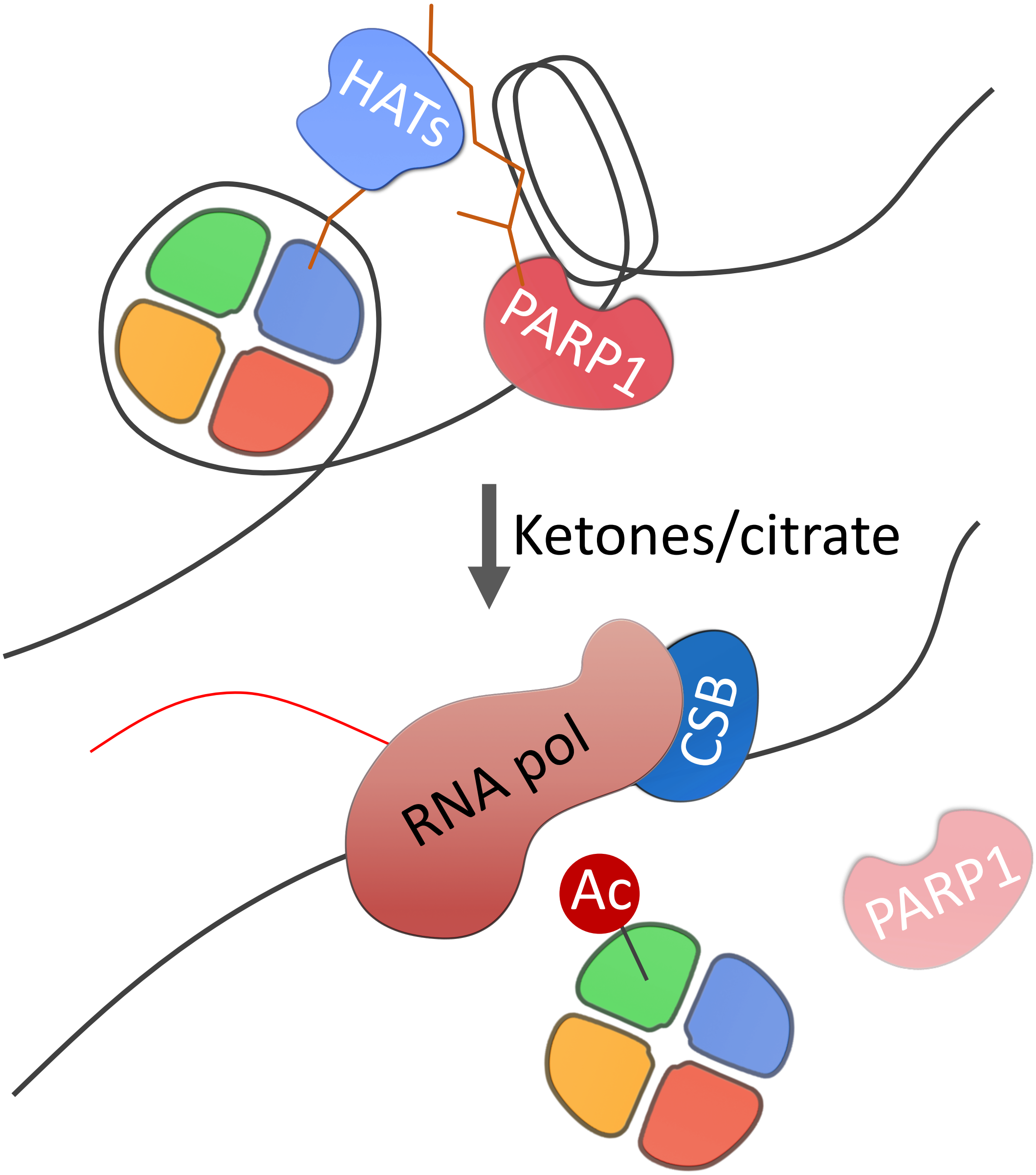


Figure S8
